# Differential growth rates and *in vitro* drug susceptibility to currently used drugs for multiple isolates of *Naegleria fowleri*

**DOI:** 10.1101/2021.10.12.464118

**Authors:** A. Cassiopeia Russell, Dennis E. Kyle

## Abstract

The free-living amoeba, *Naegleria fowleri*, which typically dwells within warm, freshwater environments, can opportunistically cause Primary Amoebic Meningoencephalitis (PAM), a disease with a mortality rate of >98%, even with the administration of the best available drug regimens. The lack of positive outcomes for PAM has prompted a push for the discovery and development of more effective therapeutics, but most studies only utilize one or two clinical isolates in their drug discovery assays. The inability to assess possible heterogenic responses to drugs among isolates from varying geographical regions hinders progress in the field due to a lack of proven universal efficacy for novel therapeutics. Herein we conducted drug efficacy and growth rate determinations for 11 different clinical isolates, including one obtained from a successful treatment outcome, by applying a previously developed CellTiter-Glo 2.0 screening technique and flow cytometry. We found some significant differences in the susceptibility of these isolates to 7 of 8 different drugs tested, all of which comprise the cocktail that is recommended to physicians by the Centers for Disease Control. We also discovered significant variances in growth rates among isolates which draws attention to the dissidence among the amoebae populations collected from different patients. The findings of this study reiterate the need for inclusion of additional clinical isolates of varying genotypes in drug assays and highlight the necessity for more targeted therapeutics with universal efficacy across *N. fowleri* isolates. Our data establishes a needed baseline for drug susceptibility among clinical isolates and provides a segue for future combination therapy studies as well as research related to phenotypic or genetic differences that could shed light on mechanisms of action or predispositions to specific drugs.

## 1. Introduction

The amphizoic amoeba, *Naegleria fowleri*, is a protozoan eukaryote that is normally found ubiquitously in warm, fresh water environments (1). Colloquially known as the “brain-eating amoeba”, it is the causative agent for primary amoebic meningoencephalitis (PAM), an acute brain disease that results from amoeba-contaminated water infiltrating the nasal cavity. This allows the invasive trophozoite stage to bind to and colonize the nasal epithelium, travel through the nasal mucosa, migrate along the neuro-olfactory nerves and traverse the cribriform plate to reach the olfactory bulbs in the central nervous system (CNS) where it feeds on neurons and damages brain membranes and meninges (1-4). Although the pathogenicity of the amoeba contributes to some of the damage, the intense immune response mounted by the host ultimately leads to death due to increased intracranial pressure and brain herniation resulting in pulmonary edema and cardiopulmonary arrest (5). The incubation period of PAM ranges from 2-15 days with >97% of cases resulting in death approximately 1 week after the initial appearance of symptoms (3, 6). This alarming mortality rate can be contributed to multiple factors, including incorrect/delayed diagnosis and ineffective clinical therapeutics. The US Centers for Disease Control and Prevention (CDC) recommends a regimen of chemotherapeutics that includes amphotericin B, an azole (ketoconazole, fluconazole, posaconazole or miconazole), azithromycin, rifampin, and miltefosine (2). More detailed guidance for healthcare providers regarding treatment can be found at the following website provided by the CDC: https://www.cdc.gov/parasites/naegleria/treatment-hcp.html.

Out of the hundreds of cases of PAM reported by the CDC, there have only been 7 survivors with confirmed diagnoses (7). All of these survivors were treated with amphotericin B, and most also received an azole, rifampin, azithromycin and miltefosine (7). Taravaud et al. have compiled case reports in an extensive literature review to show trends in treatment selections and report that amphotericin B and rifampicin are used most frequently, with fluconazole, azithromycin, and miltefosine trailing close behind (8). The frequent use of these drugs compounded with the lack of successful outcomes leads us to speculate that there are other factors at play among amoebae populations that could be contributing to treatment failure. Among these unknown factors are possible differences in growth rates that could accelerate or slow the progression of the disease and/or metabolism that could potentially conduce to differences in susceptibility to drugs among different amoeba isolates.

There are numerous studies that report the susceptibility of 1 to 3 clinical isolates of *N. fowleri* to a variety of drugs (9-28) and only 4 studies that use 5 or more different isolates (29-32). As such, we selected amphotericin B, fluconazole, ketoconazole, miconazole, posaconazole, azithromycin, rifampin, and miltefosine to assess their efficacy and determine the consistency in the response among *N. fowleri* clinical isolates of various genotypes, geographical locales, and temporal origins. Additionally, studies that show the growth rates of various isolates of this parasite have not been performed since the 80’s in which Haight et al. show that considerable variation of growth occurs for different strains of *N. fowleri* even when cultured in same medium under the same conditions (33). To characterize these variations in growth rates and also to rule out the possibility of varying susceptibility due to differential growth rates, we utilized flow cytometry to determine the doubling time of each of the strains. Henceforth, we report the results of trophocidal assays using each of the aforementioned drugs, as well as the calculated growth rates for 11 clinical isolates of *N. fowleri*.

## 2. Materials and Methods

### 2.1 Naegleria fowleri *Clinical Isolates*

*Nf69* (ATCC 30215), a clinical isolate used as a reference strain in these studies was obtained from a 9-year-old boy in Adelaide, Australia who died in 1969 (34-36), and *6088* (ATCC 30896), obtained from a 9-year-old girl in California who survived in 1978 (37, 38), were purchased from the American Type Culture Collection (ATCC). *V067*, isolated from a 30-year-old male in Arizona who died in 1987 (39, 40), *V206*, isolated from a man in Mexico who died in 1990 (39), *V413*, isolated from a 17-year-old boy in Texas who died in 1998 (39, 40), *V597*, isolated from a 10-year-old boy in Florida who died in 2007, *V631*, isolated from a 28-year-old man in in Louisiana who died in 2011 (41), *Davis*, isolated from an individual in Florida who died in 1998 (30), *HB1*, isolated from a 16-year-old boy in Florida who died in 1966 (25, 42-44), *HB4*, isolated from a female in Virginia who died in 1977 (38, 45, 46), and *Ty*, obtained from a 14-year-old boy in Virginia who died in 1969 (30, 32, 38, 47, 48) were all kindly provided by Dr. Ibne Ali at the Centers for Disease Control and Prevention (CDC).

### 2.2 Culturing of Amoebae

Trophozoites were grown axenically at 34 °C, 5% CO_2_ in non-vented 75 cm^2^ tissue culture flasks (Olympus, El Cajon, CA, USA) with Nelson’s complete medium (NCM) supplemented with 10% fetal bovine serum (FBS) and 100 units/mL penicillin and 100 mg/mL streptomycin until 80-90% confluent. For sub-culturing, flasks were placed on ice for approximately 15 minutes to detach adherent cells, and cells were collected via centrifugation at 4°C for 5 minutes at 4000 RPM. The resulting amoeba pellet was then resuspended in 1mL of NCM supplemented with 10% FBS and either passaged to new flasks or diluted further for manual counting via hemocytometer. These counts were performed in duplicate, and the resulting mean was used to calculate the dilutions needed for microwell plates (4000 cells per well) and microcentrifuge tubes (35,000 cells per tube).

### 2.3 Drug Sources and Preparations

Amphotericin B (CAS# 1397-89-3), Azithromycin (CAS# 117772-70-0), Ketoconazole (CAS# 65277-42-1), and Posaconazole (CAS# 171228-49-2) were purchased from Sigma Aldrich (St. Louis, MO). Miconazole (CAS# 22916-47-8) was purchased from Millipore Sigma (Sigma Aldrich, St. Louis, MO). Miltefosine (CAS# 58066-85-6) was purchased from Cayman Chemical (Ann Arbor, MI), rifampicin (CAS# 13292-46-1) was obtained from Fischer Scientific (Hampton, NH) and fluconazole (CAS# 86386-73-4) was purchased from the United States Pharmacopeia (Rockville, MD). Working stocks for all compounds were created by dissolving the drugs in dimethylsulfoxide (DMSO) at 5mg/mL concentrations with the exception of fluconazole, which was prepared at 25mg/mL.

### 2.4 CellTiter-Glo 2.0 in vitro Drug Susceptibility Assay

We used the CellTiter-Glo 2.0 reagent kit (CTG; Promega, Madison, WI) to determine the 50% inhibitory concentrations (IC_50_s) for each drug and amoeba isolate as previously described (17, 23, 31). To identify the optimal concentration of cells to seed for this assay and also to confirm that this falls within the linear range of the luminescence-based assay, the optimal seeding density of each of the isolates was determined by seeding a dilution series of trophozoites (ranging from 1000 cells to 10,000 cells) in Thermo Scientific Nunc Flat-bottomed 96-well MicroWell white polystyrene plates (Thermo Fisher Scientific, Waltham, MA). Drug susceptibility assays involved serial dilutions of each drug initially from 50μg/mL to 5ng/mL (250 μg/mL to 25ng/mL for fluconazole and starting at lower concentrations as needed for more potent drugs) with 4000 amoebae/well to generate dose-response curves. Control wells were cultured in triplicate per plate with 1% DMSO for the negative controls and 54.1 μ M of amphotericin B for the positive controls. Each isolate was normalized to its own controls due to variability in average luminescence produced among isolates. Plates were incubated at 34 °C, 5% CO_2_ for 72 hours.

At the 72-hour timepoint, 25uL of CellTiter-Glo 2.0 reagent was added to each well. Plates were protected from light and well contents were mixed to facilitate cell lysis via an orbital shaker at 300 RPM for 2 minutes at room temperature. Plates were equilibrated for 10 minutes, per manufacturer recommendation, to stabilize the luminescent signal prior to measurement. The luminescent signal, reported as relative light units (RLUs), is directly proportional to the amount of adenosine triphosphate (ATP) in each well and was measured at 490 nm with a SpectraMax I3X plate reader (Molecular Devices, Sunnyvale, CA, USA). To generate drug susceptibility curves, ATP RLU curve fitting was performed with Collaborative Drug Discovery Inc. software (CDD Vault, Burlingame, CA, USA). To measure percent inhibition, negative controls were calculated using 1% DMSO and positive controls were calculated with 54.1 μM amphotericin B and these controls were applied for each isolate within each plate to normalize the data. Nonlinear regressions were performed using the Levenberg–Marquardt algorithm within CDD vault. IC_50_ values were obtained from 3 biological replicates, each consisting of 2 technical replicates per drug concentration, and with the standard error of the mean (SEM). Dose response graphs were prepared by using GraphPad Prism software version 9.0 (La Jolla, CA, USA).

### 2.5 Growth Rate Calculations via Flow Cytometry

With the goal of obtaining absolute counts of the amoebae in an easily reproducible manner, we seeded 35,000 amoebae in 1.5mL of NCM supplemented with 10% FBS and 100 units/mL penicillin and 100 mg/mL streptomycin in 1.5mL microcentrifuge tubes and incubated these at 34C, 5% CO_2_ for 72h. At the 72h timepoint, tubes were centrifuged at 14,000 RPM for 5 minutes at room temperature, media was carefully aspirated from pellets and 250uL 1X PBS was added to each tube. To fix the amoebae, 250uL of 4% paraformaldehyde (PFA) was added and the pellets were resuspended via gentle pipetting. Cells were incubated in the resulting 2% PFA mixture for 15 minutes at room temperature before being centrifuged at 14,000 RPM for 5 minutes followed by careful removal of the supernatant and resuspension of the pellet in 0.5mL of 1X PBS. No viability dye was added as we wanted to limit the amount of manipulation to prevent possible loss of cells and attain more accurate counts of amoebae. An additional manual count of each of the isolates was performed in duplicate for one of the 3 biological replicates following the 72-hour incubation within microcentrifuge tubes; these data reconfirmed the accuracy of flow cytometry gating and counts (data not shown).

All samples were analyzed using a NovoCyte Quanteon flow cytometer (ACEA Biosciences, San Diego, CA, USA) to perform an absolute count based upon light scatter. The strategy used for gating cells (shown in Fig. S2) involved exclusion of debris in the reference sample (*Nf69*) as well as inclusion of the majority of events in order to account for differences in size among amoebae populations. This gating was then applied to the remaining samples to maintain consistency between isolates. This data was analyzed using FCS Express software version 7.06.0015 (De Novo Software; Pasadena, CA).

To calculate the generation time of each of the clinical isolates, the following equation for bacterial growth by binary fission was adapted to suit our needs.

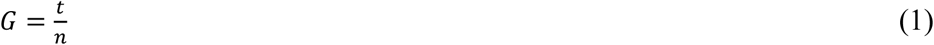

The generation time, *G*, is equal to time in hours, *t*, divided by number of generations, *n*, with *n* being the number of times the cell population doubles during the specified time interval, calculated with the following equation:

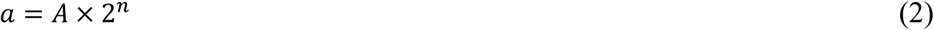

The number of amoebae at the end of the time interval was defined as *a* and the number of amoebae at the beginning of a time interval was defined as *A*. Simplification of (2) to solve for *n* and substitution into (1) provided the final equation (3) which was utilized for the reported generation time calculations:

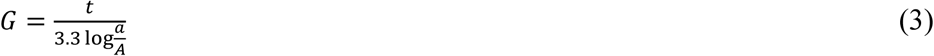

### 2.6 Genotype Determination and Method Comparison

Because the genotype for *Nf69* has not been previously reported in the literature, we used the techniques described by De Jonkheere et al (49) to determine the genotype for this isolate. Genomic DNA extractions were performed upon 5 million trophozoites of *Nf69* using Quick-DNA Miniprep Plus Kit according to manufacturer recommendations (Zymo Research, Irvine, CA, USA). In short, we utilized the following ITS primers for *Naegleria* sp.: ITSFWD (5’-AACCTGCGTAGGGATCATTT-3’), and ITSREV (5’-TTTCCTCCCCTTATTAATAT-3’). The PCR amplification was carried out with Phusion High-Fidelity PCR Master Mix with HF Buffer in 50uL reactions with 20ng of genomic DNA (New England Biolabs Inc., Ipswich, MA, USA). Reactions were run in an Agilent SureCycler 8800 (Agilent Technologies, Santa Clara, CA, USA) for 6 minutes at 94°C to allow for DNA denaturation, and then 35 cycles of 94°C for 1 minute, 55°C for 1 minute and 30 seconds, 72°C for 2 minutes with a final elongation step for 10 minutes at 72°C. Following the PCR cycling, 12uL of the PCR reaction was mixed with 2uL of 6X DNA Loading Dye (Thermo Fisher Scientific, Waltham, MA, USA) and this was run in a 1.5% agarose gel at 125V for ∼90 minutes. The GeneRuler 1kb plus DNA Ladder (Thermo Fisher Scientific, Waltham, MA, USA) was used as a size marker and the band of ∼400bp was excised and purified using the MinElute Gel Extraction Kit according to the manufacturer’s protocols (Qiagen, Hilden, Germany). The resulting purified PCR products were sent for Sanger sequencing in both the forward and reverse directions via Genewiz sequencing services (South Plainfield, NJ, US). Poor quality bases were trimmed from either end using Geneious Prime software version 2020.2.5 (Biomatters Ltd, Auckland, New Zealand), and the consensus between the forward and reverse sequences as well as duplicate samples was extracted for further analyses.

To compare genotype determination for each of the two methods that are reported in the literature (39, 49), we accessed the genome sequences recently published under BioProject PRJNA642022 and imported them to Geneious Prime software as paired reads in order to obtain pairwise alignments (options: Illumina, Paired end (inward pointing), pairs of files, 500bp) for each clinical isolate included in this study (50). We then mapped the reads to the ITSFWD and ITSREV primers using the Geneious mapper with Medium-Low Sensitivity allowing for iterations up to 5 times for fine tuning (all other options were set to default). The results of this mapping step were then *de novo* assembled to each other with the Geneious assembler (with Medium Sensitivity and default parameters; the option to dissolve the contigs and reassemble was not selected), which resulted in 2 contigs—one that bound the ITSFWD primer and one that bound the ITSREV primer. The overlapping consensus sequences from the FWD contig as well as the reverse-complement of the REV contig were extracted and genotyping was performed and reported in Table 1 as described in the results section (49).

**TABLE 1.** Clinical isolate patient/regional information, growth rates, genotypes and literature references.

| Isolate Description<br>(Alternative Identifiers) | Sex/Location/Year | Mean Growth<br>Rate (H) ± SEM * | Genotype <sup>†</sup> | References |
| --- | --- | --- | --- | --- |
| **Nf69**<br>(ATCC 30215) | M/Australia/1969 | 20.3 ± 2.1 | IV * / 5 * | 34-36 |
| **V067**<br>(V523) | M/Arizona/1987 | 21.4 ± 1.0 | III / 3 * | 39, 40, 49 |
| **V206** | M/Mexico/1990 | 24.4 ± 0.2 | I / 2 * | 39, 49 |
| **V413** | M/Texas/1998 | 18.5 ± 0.5 | I / 2 * | 39, 40, 49 |
| **V597** | M/Florida/2007 | 29.6 ± 0.7 | I / 2 * | 49 |
| **V631** | M/Louisiana/2011 | 24.6 ± 1.0 | I / 2 * | 41, 49 |
| **6088**<br>(ATCC 30896; CAMP) | F/California/1978 | 23.7 ± 0.6 | II / 1 * | 37, 38, 49 |
| **Davis**<br>(V414) | M/Florida/1998 | 22.4 ± 0.7 | I / 2 * | 30, 49 |
| **HB1**<br>(Hb) | M/Florida/1966 | 19.0 ± 1.2 | III / 3 * | 25, 42-44, 49 |
| **HB4** | F/Virginia/1977 | 21.7 ± 1.2 | III / 3 * | 38, 45, 46, 49 |
| **TY** | M/Virginia/1969 | 22.9 ± 1.9 | III / 3 * | 30, 32, 38, 47, 48, 49 |

<sup>†</sup> Genotyping reported according to 2 different methods listed in table as: Zhou et al. [Roman numerals] / De Jonckheere [Arabic numerals] (39, 49)
\* Mean growth rates and/or genotype determination made in this study

### 2.7 Statistical Analyses

We used the Z’ factor as a statistical measurement to confirm the validity of the drug susceptibility screening assay. By considering the mean and standard deviation for the positive and negative controls of each drug plate, Z’ assesses data quality and robustness of each plate to indicate the probability of false positives or negatives. All of the plates screened had excellent Z’-scores > 0.56.

To determine if there were statistically significant differences in IC_50_ for each drug and growth rates among the isolates, one-way ANOVAs with Dunnet’s multiple comparisons and Tukey-Kramer’s multiple comparisons post hoc test, respectively, were performed using GraphPad Prism version 9.1.2 (La Jolla, CA, USA).

## 3. Results

### 3.1 Comparison of drug susceptibility data among clinical isolates

We first aimed to determine the susceptibility of each clinical isolate of *N. fowleri* to 8 drugs commonly used for treatment of PAM. The IC_50_ for each drug was determined individually with three biological replicates for all 11 clinical isolates (Table 1, Table S1). We used the *Nf69* isolate as the reference for comparisons because it has been used extensively in recent drug discovery research (17, 23, 31, 51-54). As shown in Figure 1A, there was no statistically significant difference in the susceptibility of the isolates for amphotericin B when compared to the reference isolate as determined by one-way ANOVA (F(10,22)=1.711, p=0.1411). For rifampicin (Fig. 1H), although the one-way ANOVA results indicated that there was significant variance among the set of isolates as a whole, post hoc Dunnet’s multiple comparison tests showed no statistically significant differences between *Nf69* and the other 10 isolates (F(10,22)=4.618, p=0.0013). For azithromycin (Fig. 1B), the IC_50_ for *Ty* was more than four times higher than that of *Nf69*, 0.14 ± 0.05 μM versus 0.029 ± 0.001 μM, respectively. This difference was determined to be statistically significant with one-way ANOVA (F(10, 22)=2.944, p=0.0166) and post hoc Dunnett’s multiple comparisons test (p=0.0066). When evaluating the effectiveness of fluconazole (Fig. 1C), we utilized an upper cutoff of 820 μM and 5 of the 11 isolates (including the reference) surpassed this cutoff, showing a lack of inhibition by this compound in nearly half of the clinical isolates tested. Furthermore, the IC_50_ values for the remaining isolates were found to be significantly lower than *Nf69*, which was reported as 820 μM due to a lack of inhibition even at the highest concentration of fluconazole tested (F(10,12)=34.91, p<0.0001). For *V067, V206, V597, V631, HB1*, and *Ty*, the IC_50_s for fluconazole were 65.3 ± 22.7 μM, 109 ± 16.0 μM, 196.7 ± 36.7 μM, 216.7 ± 80.9 μM, 7.73 ± 0.87 μM, and 150.7 ± 52.3 μM, respectively.

**FIGURE 1.**
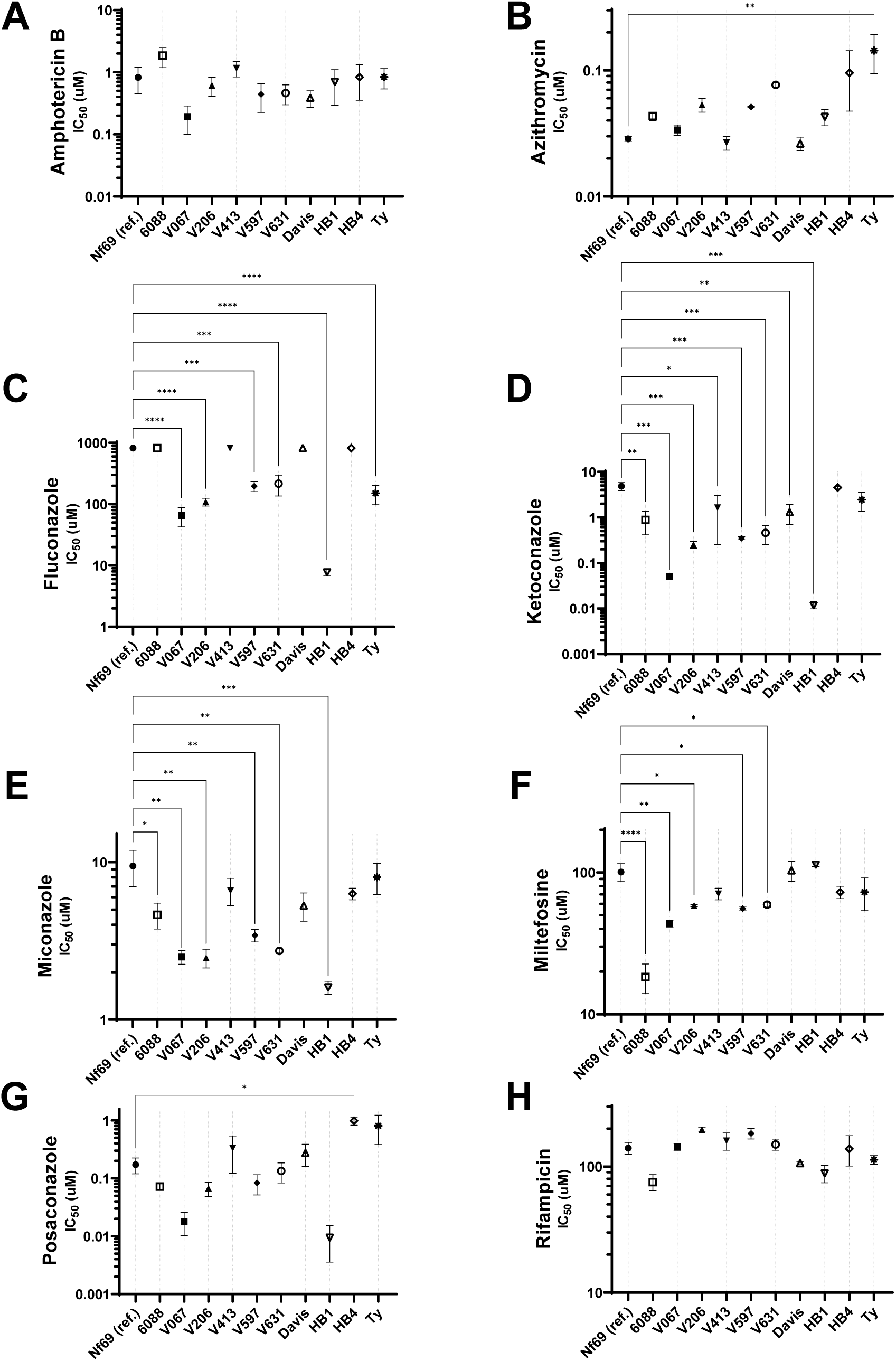
The IC_50_s of the 11 isolates determined using the CellTiter-Glo 2.0 72h assay are shown for each of the tested drugs: amphotericin B **(A)**, azithromycin **(B)**, fluconazole **(C)**, ketoconazole **(D)**, miconazole **(E)**, miltefosine **(F)**, posaconazole **(G)**, and rifampicin **(H)**. Statistical significance was determined by one-way ANOVA with *Nf69* as the reference strain. ‘*’ = Significant (0.01 < p < 0.05), ‘**’ = Very significant (0.001 < p < 0.01), ‘***’ = Extremely significant (0.0001 < p < 0.001), ‘****’ = Extremely significant (p < 0.0001).

Eight of the clinical isolates showed significantly increased susceptibility to ketoconazole when compared to *Nf69* which had an IC_50_ of 4.87 ± 0.97 μM (F(10,22)=6.913, p<0.0001). The IC_50_ values were 0.89 ± 0.47 μM, 0.05 ± 0.006 μM, 0.25 ± 0.043 μM, 1.64 ± 1.38 μM, 0.35 ± 0.026 μM, 0.46 ± 0.21 μM, 1.31 ± 0.62 μM, and 0.012 ± 0.001 μM for *6088, V067, V206, V413, V597, V631, Davis*, and *HB1* respectively. For miconazole (Fig. 1E), 6 of the isolates were significantly more susceptible than *Nf69* (F(10,22)=5.338, p=0.0005), with IC_50_s of 9.48 ± 2.4 μM, 4.63 ± 0.86 μM, 2.5 ± 0.34 μM, 2.47 ± 0.34 μM, 3.43 ± 0.32 μM, 2.73 ± 0.088 μM, and 1.6 ± 0.15 μM for *Nf69, 6088, V067, V206, V597, V631*, and *HB1* respectively. Five isolates showed increased susceptibility to miltefosine compared to the value of 100.7 ± 14.6 μ M for *Nf69* (F(10,22)=8.64, p<0.0001), with IC_50_s of 18.3 ± 4.3 μM, 43.7 ± 2.4 μM, 58.3 ± 0.88 μM, 55.7 ± 2.0 μM, and 59.3 ± 2.3 μM for *6088, V067, V206, V597* and *V631* respectively. Lastly, *HB4* was the only isolate to show a significant decrease in susceptibility to posaconazole with an IC_50_ of 0.98 ± 0.16 μM (Fig. 1G), which is more than 4 times higher than that of 0.17 ± 0.052 μM for *Nf69* (F(10,22)=4.449, p=0.0017). It is possible that sulfisoxazole, one of the drugs administered to the surviving patient in 1978 (*6088*), could have contributed to this positive outcome, so we also tested its efficacy on each of the isolates. Although this compound was tested at concentrations up to 940 μM, it was not found to affect the growth of any of the isolates when compared to controls (data not shown). Additional IC_50_ values (± SEM) can be found in Table S1.

### 3.2 Growth Rate Results

By utilizing the absolute count feature of FCS Express software when analyzing our flow cytometry data, we were able to develop a semi-automated counting technique for amoebae that excluded debris that are common when culturing free-living amoebae; this method was used to monitor the growth rates of the 11 clinical isolates included in this study. Shown in Table 1, the generation time of these amoebae ranged from 18.5 ± 0.5 hours for *V413* to 29.6 ± 0.7 hours for *V597*, which is about 1.5 times slower than the fastest growing isolate. The reference isolate, *Nf69*, was one of the fastest growing isolates with a generation time of 20.3 ± 2.1 hours. The overall average growth rate of the clinical isolates was found to be approximately 22.4 ± 0.9 hours. We found multiple statistically significant differences among the isolates as determined by one-way ANOVA (F (10, 22) = 7.184, p<0.0001). Shown in Table S2, post hoc Tukey-Kramer’s multiple comparisons test showed significant differences between the generation times of *Nf69* vs. *V597* (p=0.0004), *V067* vs. *V597* (p=0.0019), *V206* vs. *V413* (p=0.0445), *V413* vs. *V597* (p<0.0001), *V413* vs. *V631* (p=0.0343), *V597* vs. *6088* (p=0.0445), *V597* vs. *Davis* (p=0.0076), *V597* vs. *HB1* (p<0.0001), *V597* vs. *HB4* (p=0.0028), and *V597* vs. *TY* (p=0.0152). *V597*, which was the slowest growing strain, accounted for 8 of the 10 differences in generation times while *V413—*the fastest growing strain—accounted for the other 2 differences. To confirm that the growth rate was not a confounding factor when comparing susceptibility of drugs among isolates, we performed a bivariate analysis of the growth rates versus the IC_50_s (Fig. S3) and found that there was no correlation in the susceptibility of the amoebae to the drugs when compared to the speed of their growth.

### 3.3 Nf69 Genotyping Results

In addition, we wanted to compare the drug susceptibility of the clinical isolates with their genotype. There is published ITS genotype information, determined based upon the techniques described by Zhou et al., for all of the isolates except *Nf69* (39, 50). Following the attainment of Sanger sequencing results for the PCR products for *Nf69*, we trimmed off poor quality base pairs and aligned the sequences obtained for the forward primer sequencing samples as well as the reverse primer sequencing samples to attain a 446 bp sequence (GenBank accession MZ494674). We identified the forward primer and the complement of the reverse primer within the sequence, and then followed the recommendation to make a genotype determination (49). We identified 1 copy of “repeat 2” (ATGGTAAAAAAGGTGAAAACCTTTTTTT), 1 copy of “main 2” (ATGGTAAAAAAGGTGAAAACCTTTTTT), as well as 1 copy of “main 1” (CCATTTACAAAAAAT). To make the final identification, we found “repeat 1” (ATGGTAAAAAAGGTGT) to have a 2 base-pair deletion of the A and G residues at the 11^th^ and 12^th^ positions, respectively, and also a C nucleotide at position 31 in the 5.8S rDNA that followed the 84 base-pair ITS1 sequence. With all of these findings, we were able to conclude that *Nf69* falls within the genotype 5 category previously described in the literature for the *N. fowleri* species (49, 55). We also followed the less detailed protocol described by Zhou et al. and found that *Nf69* falls within the genotype IV category (39).

A recent study by Joseph et al. sequenced the genomes for each of the isolates tested in our study and we downloaded and processed this data in order to perform the aforementioned genotyping technique and compare the genotype results attained with the system created by De Jonckheere with those reported based upon Zhou et al. (39, 49, 50). Using Geneious Prime software, we pairwise aligned the forward and reverse reads for each clinical isolate and then mapped these to each of the 2 ITS primers. We then *de novo* assembled these reads which resulted in 2 contigs per isolate, one which bound the FWD primer and the other the REV primer. We extracted and aligned sequences from each contig to obtain a consensus sequence that was then processed to determine the genotypes (see Materials and Methods for a more detailed description). Shown in Table 1, we found that the isolates defined at genotype I by Zhou et al.’s guidelines are defined as genotype 2 by De Jonckheere’s methods, and those that are genotype II are genotype 1, respectively (39, 49). Additionally, genotype III was equivalent to genotype 3, and genotype IV according to Zhou et al. was genotype 5 by De Jonckheere’s definition.

## 4. Discussion

This is the first study showing the *in vitro* susceptibility of more than 10 clinical isolates of *Naegleria fowleri* to the currently recommended chemotherapeutics for the devastating neurological infection, PAM. In the literature, the *in vitro* drug susceptibility testing for PAM has progressed incrementally since the disease was first discovered in the late 1960’s. Initial assays developed for amoebae viability to chemotherapeutics all involved manual counting and potentially subjective phenotypic determination. In 1976, Duma et al. performed a study with 6 different clinical isolates of *N. fowleri* and tested amphotericin B, clotrimazole and miconazole *in vitro* against all of the organisms (32). Of the clinical isolates tested, *TY* is the only one that we also utilized in our study and upon comparing our data for amphotericin B with their MIC of 0.422μM, we found that it overlaps nicely with our IC_50_ of 0.73 ± 0.3μM (shown in Supp. Table 1). Additionally, they reported a MIC of 2.98μM for miconazole against *TY*, which is similar to the IC_50_ of 7.6 ± 1.8μM in this study (32).

Subsequent studies in the late 1970s and early 1980s all reverted to the usage of a single clinical isolate to determine chemotherapeutic susceptibility *in vitro* (9-13). The practice of utilizing multiple isolates to reconfirm propensities to novel therapeutics was not revisited until 2006 when Schuster et al. utilized 3 different strains to perform *in vitro* testing of miltefosine and voriconazole (27). There was no overlap in the isolates used for this study compared to ours, however miltefosine was used and the authors showed that a minimum of 55μM was required to attain amoebicidal conditions, which falls within our range of IC_90_s from 21.6μM to >120μM shown across isolates (also see Supp. Table 1). Importantly, these authors also observed strain variations in sensitivity to the drugs they tested on amoebae (27). Further technical advances were made by Kim et al. in 2008 with the optimization of a colorimetric lactate dehydrogenase release assay to measure amoebicidal activity of compounds *in vitro* (28). This provided the groundwork for the development and optimization of high-throughput screening assays to identify novel chemotherapeutics to treat PAM (16, 17).

Surprisingly, even with the establishment of high-throughput assays for testing compounds against this amoeba, it took another decade for the inclusion of more than a single isolate of *N. fowleri* in drug testing studies. In 2018, Peroutka et al. used 2 different isolates, *LEE* and *HB1*, to determine amoeba viability against auranofin. The authors noted a significant difference in sensitivity between the 2 amoebae and indicated that clinical outcomes could be dependent on individual strain susceptibilities to drugs (19). In 2019, Colon et al. utilized a high-throughput screening method to identify novel drugs to treat PAM using 6 different isolates: *Nf69, V067, V206, V413, V414*, and *V596*. They compared the *in vitro* drug susceptibility of 4 chemotherapeutics and showed no statistical difference between *Nf69* and the 5 other isolates for amphotericin B, azithromycin, miltefosine, and posaconazole (31). Our study, which incorporates 4 of the 6 clinical isolates used by Colon et al., reiterates the lack of significant differences between isolates for amphotericin B as well as azithromycin and posaconazole. However, due to our inclusion of additional strains, we were able to detect variance in two of the isolates not tested by Colon et al.—*TY* for azithromycin, and *HB4* for posaconazole—which further emphasizes the need for multi-isolate drug testing in *N. fowleri*. To further showcase the recent trend in the right direction, in 2020 Escrig et al. incorporated 5 different clinical isolates of varying genotypes and geographic origins to showcase the variance in susceptibility to the tested chemical compounds (30). Their average EC_50_ of 37.8μM for miltefosine falls within the range of IC_50_s obtained in our study for the same drug. With an average EC_50_ 0.094μM for amphotericin B, their data indicated a more potent response to this molecule than ours, but this could be due to differences in assay parameters (e.g. 10,000 vs. 4,000 amoebae/well. and a 48h timepoint vs. 72h), or potential batch-to-batch variation in molecule potency.

Furthermore, we report the sequence of a portion of the SSU rRNA gene, ITS1, 5.8S rRNA gene, ITS2 as well as a portion of the LSU RNA gene of *Nf69* that we attained in order to determine the genotype of this isolate (GenBank accession #: MZ494674). We initially endeavored to ascertain the genotyping parameters by referring to the resource used by Ali et al. in their recent publication in which they defined the *TY* isolate as genotype III (48). This guideline provided by Zhou et al. which uses Roman numerals and defines genotypes I-VI (39), is outdated and the repeats and various components of the ITS1 as well as 5.8S rRNA gene are less detailed when compared to the more recent review by de Jonckheere which uses Arabic numerals to differentiate types based upon the evolutionary consideration of which type likely appeared first (49). As such, there is a need for the establishment of a universal set of genotyping parameters to maintain consistency among publications and future research when characterizing isolates of *N. fowleri*.

Not only have we shown statistically significant variability in IC_50_s among the clinical isolates that were tested, but we also calculated significant differences in growth rates by culturing each isolate in controlled equivalent conditions. Previous research by Weik et al. shows that agitated cultures allow for a speedy generation rate of approximately 5.5 hours with higher maximum yields of amoebae when compared to unagitated amoebae (56). This difference in generation time is likely attributed to the higher incubation temperature of 37° C—which is more conducive to growth of bacterial/fungal contamination and also less realistic when compared to the natural temperature of the bodies of water that the amoebae are found in, hence our selection of 34° C—and also due to the maximum population density of the small volume of media used in this study. We speculate that our calculated doubling rates would likely be lower with higher temperatures or increased volume/allotted growth area.

Upon diagnosis, the timeframe for treatment is already highly compacted due to the fulminant nature of PAM, and the use of drugs that the amoebae have varying levels of innate resistance to is potentially detrimental. This data provides valuable evidence that the universal approach of applying the recommended cocktail of drugs might not be the most effective one. Overall, a lack of variety in genotypic as well as geographic origins could lead to the premature conclusion that a newly discovered compound or scaffold is universally effective against *N. fowleri* when there could be a notable difference in activity across different isolates of the parasite. Thus, the findings in this paper draw attention to the changes that need to be made in the field in treating and discovering new therapeutics for this deadly disease. Moving forward, multiple isolates of varying genotypes should be used when determining the susceptibility of this parasite to prospective drugs and bioactive molecules. This has long been the standard of practice for antimicrobial drugs and our data support the need for this change to be adopted for drug discovery for *N. fowleri*. The scientific and medical community also needs to reevaluate the effectiveness of the currently recommended therapeutics for treating PAM. It is important to recognize the most commonly used drug regimen(s) were derived empirically and based upon the few successful treatment regimens. More recently, repurposed drugs (e.g., miltefosine) have emerged from laboratory studies, yet the entire drug cocktail used to treat PAM does not have sufficient in vitro or in vivo (mouse model) data. For example, we have shown that fluconazole and rifampicin are not potent for the majority of the 11 isolates tested and thus continued use of these two drugs in the recommended treatment regimen should be reconsidered. As for azoles, we recommend treating patients with posaconazole rather than the less potent azoles fluconazole, miconazole and ketoconazole. The overarching need for this disease is a more targeted set of recommendations according to the genotype, growth rate and susceptibility of the specific isolate and the components of the therapeutic cocktail should be catered to these important elements in order to give patients a higher chance for surviving an infection by *N. fowleri*. The empirical results reported herein should be considered in light of our reported efficacy of single-entity therapy versus the standard combinational therapy for PAM. Future studies that determine combinational effects of the different drugs, whether they be synergistic or antagonistic to each other, should be performed on multiple isolates. Performing these studies with the combination of 3 or more drugs might be challenging but is warranted due to the acute nature of this infection and the urgent need for more effective treatment options.

## 5. Conclusion

In conclusion, we determined that there is a statistically significant difference in the susceptibility to the majority of the currently recommended drugs for PAM among clinical isolates of *N. fowleri*. This data shows that the current therapeutic recommendations should be re-examined and helps to establish a baseline for drug susceptibility among different clinical isolates. It also paves the way for the identification of differences between isolates, whether they are genetic elements such as SNPs or other mutations, or phenotypic elements that may provide a hint toward predisposition to specific drugs. Regardless of the approach, we show that a more targeted methodology is what is needed to focus on the specific amoebae infecting a patient and subsequently increase the odds for survival.

## 6. Data Availability

The *Naegleria fowleri ‘Nf69’* 446 bp sequence containing a portion of the SSU rRNA, ITS1, 5.8S rRNA, ITS2, and a portion of the LSU rRNA has been deposited to GenBank under accession number MZ494674, https://www.ncbi.nlm.nih.gov/nuccore/MZ494674.

## Notes

### Competing Interest Statement

The authors have declared no competing interest.

