## Supplemental Material for "Differential growth rates and *in vitro* drug susceptibility to currently used drugs for multiple isolates of *Naegleria fowleri*"

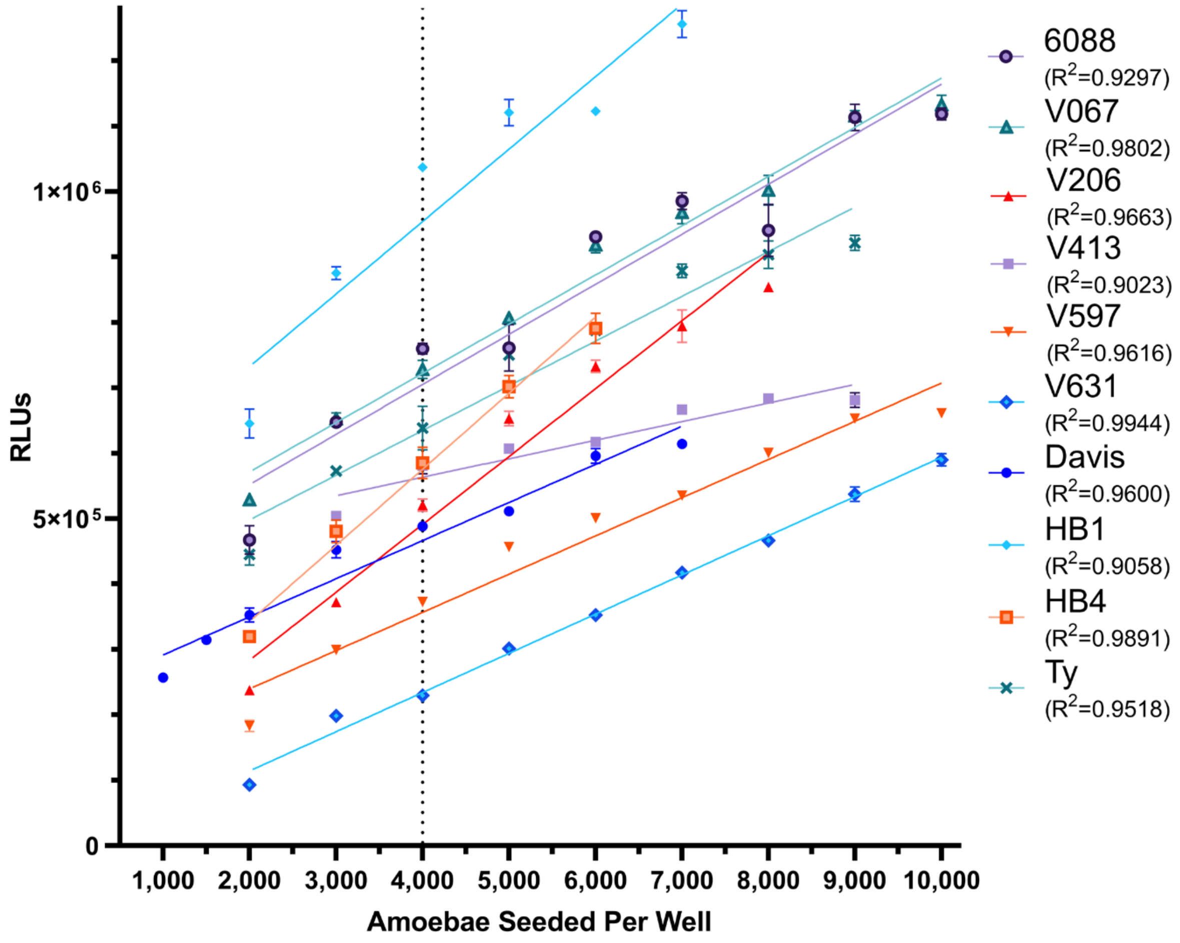


**FIG S1** Optimal seeding density for the CellTiter Glo 2.0 kit determined by linear regression with at least 5 serial dilutions per clinical isolate. Three technical replicates were performed per concentration per clinical isolate. The optimal seeding density for *Nf69* was previously determined to be 4000 amoebae per well (1).

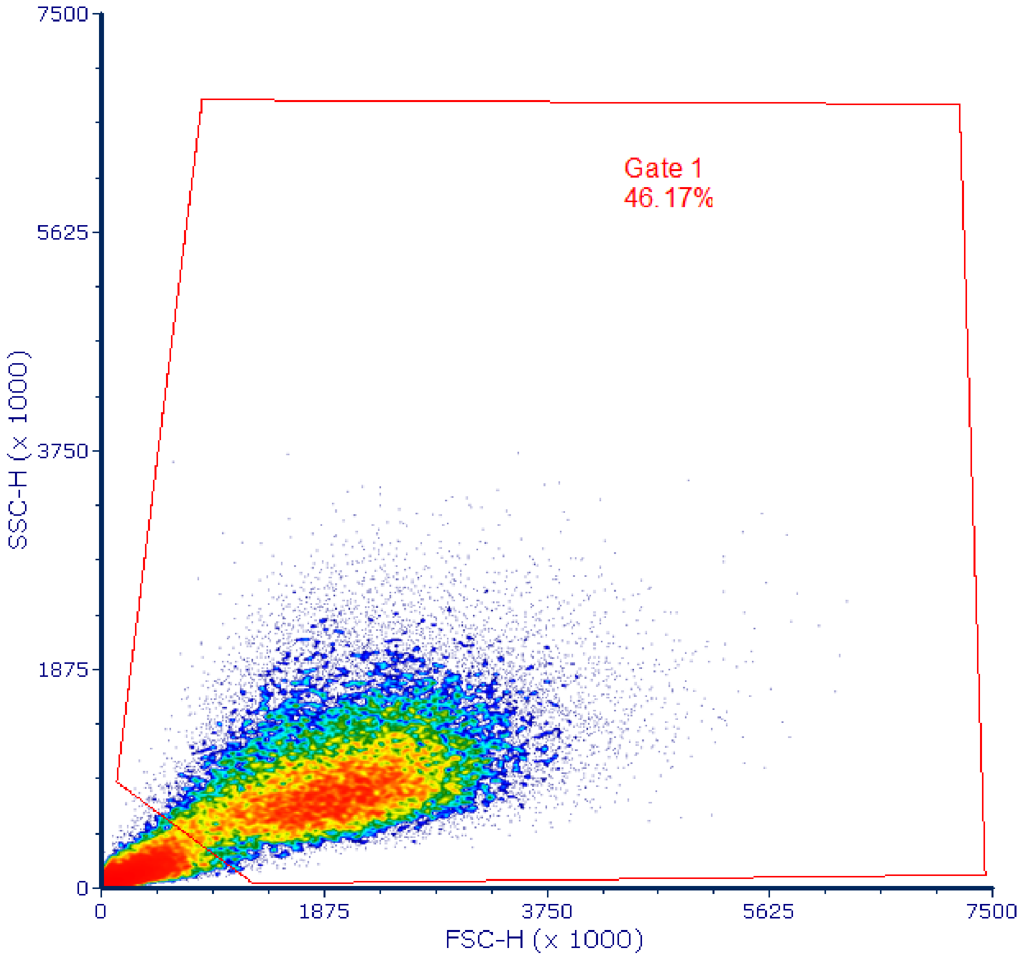


**FIG S2** Representative gating strategy in a reference sample (*Nf69*) utilized for the forward-scatter (FSC-H) versus side-scatter (SSC-H) results attained for all clinical isolates. This gating strategy was applied to all samples in order to maintain consistency in the absolute counting of amoebae.

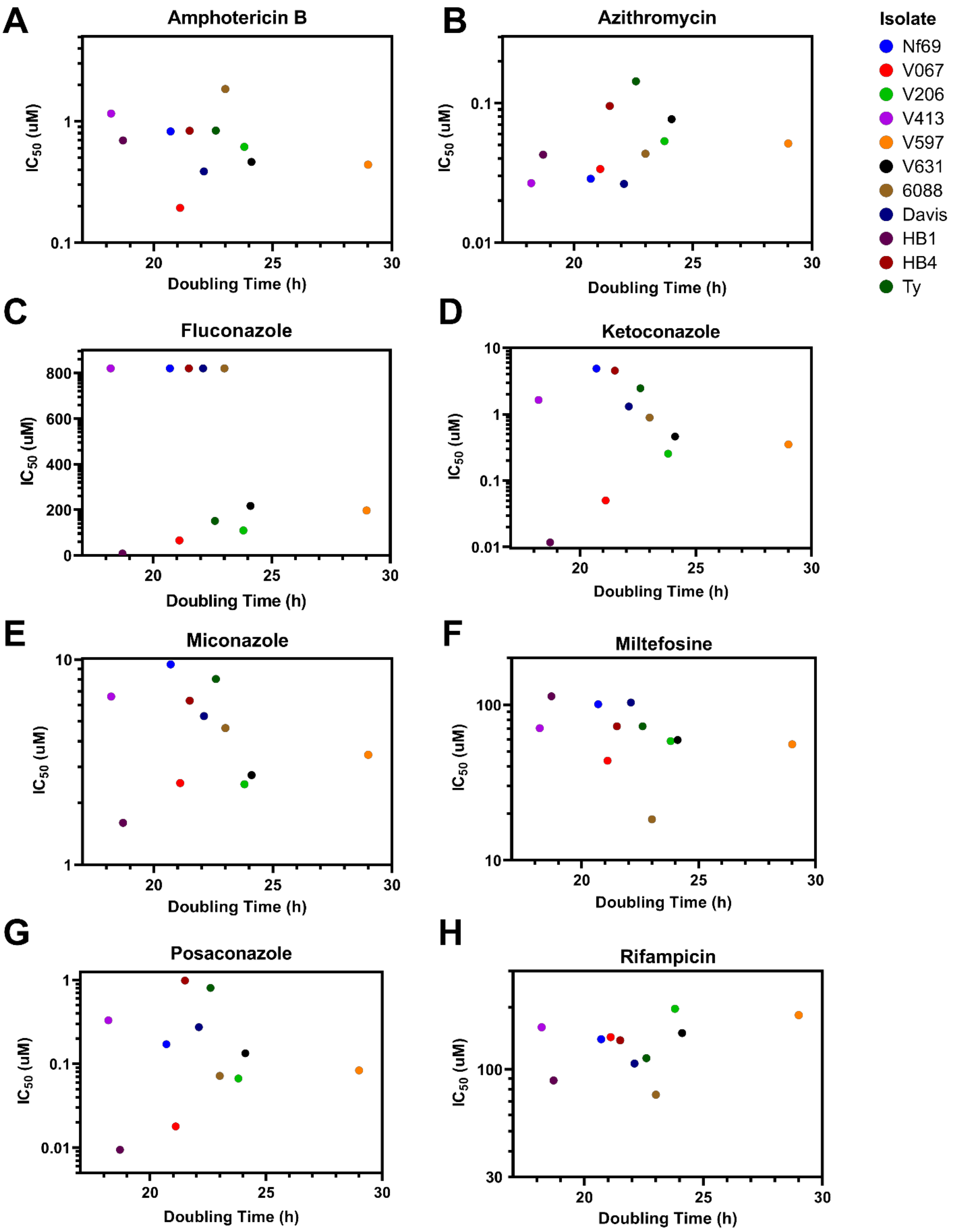


**FIG S3** Bivariate analysis of doubling time vs IC50 for each of the drugs and isolates. These data suggest there is no correlation between doubling time and susceptibility to any of the drugs tested.

**TABLE S1** IC50/90s (µM) ± the standard error of the mean determined for 11 clinical isolates of *N. fowleri*. We used 3 biological replicates, each with 2 technical replicates per concentration tested to generate dose response data.

| Clinical Isolate | | Chemotherapeutic | | | | | | | |
|---|---|---|---|---|---|---|---|---|---|
| | | Amphotericin B | Azithromycin | Fluconazole | Ketoconazole | Miconazole | Miltefosine | Posaconazole | Rifampicin |
| Nf69 | $IC_{50}$ ± SEM | 0.63 ± 0.37 | 0.026 ± 0.001 | > 820* | 4.69 ± 0.97 | 8.84 ± 2.45 | 98.3 ± 14.6 | 0.16 ± 0.05 | 138.4 ± 15.3 |
| | $IC_{90}$ ± SEM | 0.69 ± 0.39 | 0.028 ± 0.000 | > 820* | 10.3 ± 1.52 | 12.7 ± 2.05 | 116.0 ± 12.3 | 0.36 ± 0.08 | >300* |
| 6088 | $IC_{50}$ ± SEM | 1.61 ± 0.66 | 0.04 ± 0.003 | > 820* | 0.64 ± 0.47 | 4.48 ± 0.87 | 18.6 ± 3.93 | 0.07 ± 0.01 | 73.9 ± 10.9 |
| | $IC_{90}$ ± SEM | 1.73 ± 0.74 | 0.42 ± 0.25 | > 820* | 2.90 ± 2.15 | 7.76 ± 1.25 | 21.6 ± 4.10 | 0.57 ± 0.20 | >300* |
| V067 | $IC_{50}$ ± SEM | 0.16 ± 0.09 | 0.03 ± 0.003 | 54.2 ± 22.7 | 0.05 ± 0.01 | 2.47 ± 0.25 | 43.5 ± 2.40 | 0.01 ± 0.01 | 142.8 ± 8.82 |
| | $IC_{90}$ ± SEM | 0.16 ± 0.10 | 0.05 ± 0.004 | >820* | 0.92 ± 0.09 | 4.42 ± 0.22 | 82.9 ± 8.19 | 0.14 ± 0.10 | >300* |
| V206 | $IC_{50}$ ± SEM | 0.52 ± 0.21 | 0.05 ± 0.007 | 106.8 ± 15.9 | 0.25 ± 0.04 | 2.42 ± 0.34 | 58.3 ± 0.88 | 0.06 ± 0.02 | 196.3 ± 8.82 |
| | $IC_{90}$ ± SEM | 0.82 ± 0.44 | 0.10 ± 0.02 | >820* | 2.72 ± 0.90 | 9.04 ± 0.47 | 70.0 ± 0.58 | >0.36* | >300* |
| V413 | $IC_{50}$ ± SEM | 1.08 ± 0.32 | 0.026 ± 0.003 | > 820* | 0.66 ± 1.38 | 6.30 ± 1.31 | 70.0 ± 6.64 | 0.21 ± 0.21 | 156.4 ± 25.2 |
| | $IC_{90}$ ± SEM | 1.34 ± 0.37 | 0.033 ± 0.006 | > 820* | 1.65 ± 4.65 | 12.1 ± 4.49 | 96.5 ± 21.1 | >1.4* | >300* |
| V597 | $IC_{50}$ ± SEM | 0.35 ± 0.21 | 0.051 ± 0.0003 | 190.5 ± 36.7 | 0.35 ± 0.03 | 3.40 ± 0.32 | 55.6 ± 2.03 | 0.07 ± 0.03 | 181.6 ± 17.6 |
| | $IC_{90}$ ± SEM | 0.45 ± 0.22 | 0.054 ± 0.001 | >820* | 2.74 ± 0.48 | 8.67 ± 1.19 | 64.3 ± 0133 | >0.36* | >300* |
| V631 | $IC_{50}$ ± SEM | 0.40 ± 0.16 | 0.08 ± 0.003 | 172.7 ± 80.9 | 0.38 ± 0.21 | 2.73 ± 0.09 | 59.2 ± 2.33 | 0.11 ± 0.05 | 148.5 ± 15.3 |
| | $IC_{90}$ ± SEM | 0.52 ± 0.29 | 0.16 ± 0.04 | >820* | 6.68 ± 3.08 | 10.0 ± 1.33 | 67.0 ± 4.84 | >0.36* | >300* |
| Davis | $IC_{50}$ ± SEM | 0.35 ± 0.12 | 0.03 ± 0.003 | > 820* | 1.03 ± 0.62 | 5.08 ± 1.08 | 100.3 ± 16.7 | 0.21 ± 0.11 | 106.6 ± 3.33 |
| | $IC_{90}$ ± SEM | 0.41 ± 0.18 | 0.05 ± 0.005 | > 820* | 4.12 ± 0.87 | 8.73 ± 0.62 | >120* | >1.4* | >300* |
| HB1 | $IC_{50}$ ± SEM | 0.50 ± 0.40 | 0.04 ± 0.006 | 7.64 ± 0.87 | 0.01 ± 0.001 | 1.58 ± 0.15 | 113.2 ± 3.33 | 0.006 ± 0.006 | 85.9 ± 14.1 |
| | $IC_{90}$ ± SEM | 0.55 ± 0.46 | 0.41 ± 0.27 | 12.0 ± 3.10 | 0.02 ± 0.003 | 1.90 ± 0.06 | >120* | 0.008 ± 0.012 | >300* |
| HB4 | $IC_{50}$ ± SEM | 0.61 ± 0.48 | 0.07 ± 0.048 | > 820* | 4.51 ± 0.30 | 6.25 ± 0.53 | 72.0 ± 7.27 | 0.96 ± 0.16 | 128.9 ± 37.2 |
| | $IC_{90}$ ± SEM | 0.90 ± 1.16 | 0.70 ± 1.01 | > 820* | 11.7 ± 1.53 | 8.92 ± 0.64 | 97.7 ± 22.6 | 4.96 ± 2.97 | >300* |
| Ty | $IC_{50}$ ± SEM | 0.73 ± 0.30 | 0.12 ± 0.049 | 126.7 ± 52.3 | 1.50 ± 1.10 | 7.60 ± 1.80 | 68.3 ± 18.8 | 0.56 ± 0.42 | 112.7 ± 8.82 |
| | $IC_{90}$ ± SEM | 0.88 ± 0.48 | 2.31 ± 0.92 | >820* | 4.42 ± 2.15 | 13.5 ± 1.33 | >120* | 4.35 ± 2.06 | >300* |

* $IC_{50/90}$ values exceeded the maximum concentration of drug tested for this clinical isolate

**TABLE S2** Data from post hoc comparisons of calculated Growth Rates among the 11 isolates using Tukey-Kramer's Multiple Comparisons test. The mean difference is determined from the calculated average growth rate of 3 biological replicates per isolate.

| Comparisons (A vs. B) | Mean Difference (A-B) | 95% Confidence Interval of Difference | Significance | Multiplicity Adjusted P-value |
|---|---|---|---|---|
| Nf69 vs. V597 | -9.3 | -15.11 to -3.490 | *** | 0.0004 |
| V067 vs. V597 | -8.2 | -14.01 to -2.390 | ** | 0.0019 |
| V206 vs. V413 | 5.9 | 0.08962 to 11.71 | * | 0.0445 |
| V413 vs. V597 | -11.1 | -16.91 to -5.290 | **** | <0.0001 |
| V413 vs. V631 | -6.1 | -11.91 to -0.2896 | * | 0.0343 |
| V597 vs. 6088 | 5.9 | 0.08962 to 11.71 | * | 0.0445 |
| V597 vs. Davis | 7.2 | 1.390 to 13.01 | ** | 0.0076 |
| V597 vs. HB1 | 10.6 | 4.790 to 16.41 | **** | <0.0001 |
| V597 vs. HB4 | 7.9 | 2.090 to 13.71 | ** | 0.0028 |
| V597 vs. Ty | 6.7 | 0.8896 to 12.51 | * | 0.0152 |

'*' = Significant ($0.01 < p < 0.05$), '**' = Very significant ($0.001 < p < 0.01$), '***' = Extremely significant ($0.0001 < p < 0.001$), '****' = Extremely significant ($p < 0.0001$).
